# Optimality of the spontaneous prophage induction rate

**DOI:** 10.1101/546275

**Authors:** Michael Cortes, Jonathan Krog, Gábor Balázsi

## Abstract

Lysogens are bacterial cells that have survived after being infected by bacterial viruses called bacteriophages. Instead of being killed by the virus, the infected cell survives by integrating the viral DNA into its own genome. This is only possible with “temperate” bacteriophages which do not always lyse their host to reproduce, but sometimes replicate passively using the lysogenic pathway. After an infection resulting in lysogeny, the lysogen continues to grow and divide normally, seemingly unaffected by the integrated viral genome which is now referred to as a prophage. However, the prophage can have an impact on the host’s phenotype and overall fitness in certain environments. This makes competition between the lysogen and its nonlysogen counterpart possible because both cells have different genomes and potentially different growth rates. Additionally, the prophages within the lysogens are capable of spontaneously reverting back to the lytic pathway via spontaneous prophage induction (SPI), causing death of the lysogen and the release of new progeny phages. These new phages can then lyse or lysogenize other susceptible nonlysogens, thereby impacting the competition between lysogens and nonlysogens. In a scenario with differing growth rates, it is not clear whether SPI would be beneficial or detrimental to the lysogens since it directly causes cell death but also attacks nonlysogenic competitiors, either lysing or lysogenizing them. In this work we study the evolutionary dynamics of a mixture of lysogens and nonlysogens and derive general conditions on the rate of SPI resulting in lysogens displacing nonlysogens. We show that there exists an optimal SPI rate, and apply the model to bacteriophage *λ*. We find that the model can explain why the experimentally measured SPI rate for phage *λ* is so low. We also investigate the impact of stochasticity and conclude that even at low copy numbers the SPI rate can still be fairly low while still providing an advantage to the lysogens.

## 1 Introduction

Bacteriophages (or phages for short) are viruses which infect bacterial cells. By infecting bacteria, the phages can replicate themselves using the host’s cellular machinery. The replicated phages can then destroy the host cell and are released into the environment which is known as the lytic or lysis pathway. Lysis is the typical outcome of infection, but some phages can sometimes follow alternative pathway after infection called lysogeny. Phage *λ* is a well-known example for such phages called “temperate” [20]. In the lysogenic pathway, the phages integrate their viral DNA into the host’s genome. The host cell, now called a lysogen, is seemingly unaffected and continues to grow and divide normally. Typically this lysogenic state is very stable and the cell remains a lysogen after many cell divisions. Sometimes the lysogen can revert back to the lytic pathway in a process called prophage induction [18]. Prophage induction can be triggered by various environmental factors, such as DNA damage by UV radiation, via induction of the host’s SOS response. Prophage induction can also occur spontaneously in a process termed spontaneous prophage induction or SPI [18]. SPI is likely due to spontaneous accumulation of DNA damage initiating the host’s SOS response that occurs during cell replication [5, 16, 18, 21].

When a bacterial cell is converted to a lysogen, its genome now contains viral genes. This means that the lysogens genome is different from the nonlysogen’s genome, and as a result it is possible that natural selection can occur between the lysogens and nonlysogens. The integrated viral genes can be expressed within the lysogen and affect the lysogen’s phenotype and fitness. For many lysogenized cells, the lysogen strain has decreased sensitivity to antibiotics, increased biofilm formation, and virulence [2, 14, 15, 26, 27] due to the integrated viral genes. The additional viral genes could also slow cell growth because expression of extra viral genes could deplete host cell resources that promote cell division [22, 23]. For example, it has been shown that multiple phage *λ* infections reduce host cell transcription and translation [8, 13, 25]. Taken together, these facts imply that lysogens could outcompete nonlysogens or vice versa because of their differing growth rates or fitnesses in certain environments. Thus, in the long run the lysogens could be naturally selected over the nonlysogens or vice versa. Evidence for competitive advantage of lysogens over their nonlysogenized counterparts comes from experiments showing that mixtures of lysogens and lysogen-cured bacteria tend towards a state in which the lysogen starts to displace the lysogen-cured strain [3, 7].

In a mixture of only lysogens and nonlysogens, SPI events could cause some lysogens to lyse, thereby releasing phages into the mixture. The phages can then infect the nonlysogens either killing them via lysis or converting them into lysogens. The lysogens are not affected by additional infections because they are immune to superinfection. Thus, SPI could give the lysogens a competitive advantage, increasing the chances they will be naturally selected. On the other hand, SPI can also be costly to lysogenic fitness because SPI is concomitant with cell death. If there are no nonlysogens left to convert, SPI simply adds to the intrinsic lysogen death rate. It has been suggested that SPI enables lysogens to spread viral DNA within the bacterial population by steady lytic-killing and lysogenic-conversion of the nonlysogens [18]. However, from the perspective of the lysogen, it remains unknown precisely how high or low should the SPI rate be for the lysogens to be naturally selected over the nonlysogens? In other words, does there exist an optimal SPI rate enabling the natural selection of the lysogens over the nonlysogens? If so, then how does this optimal SPI rate depend, if at all, on ecological factors?

In this work, we use replicator dynamics and simulation [19] to gain a theoretical understanding of how SPI influences the competition between lysogens and nonlysogens under general parameter sets, and to identify an optimal SPI rate enabling the lysogens to be naturally selected over the nonlysogens, which is comparable with experimental values. Our model is intended to be general such that it can theoretically describe the competition between any lysogen and its nonlysogen counterpart so long as the lysogen can undergo SPI. We quantitatively compare our model to the lysogens from bacteriophage *λ* infecting *E. coli* because many of the model’s parameters for the phage *λ* - *E. coli* system have been measured previously [20]. Our major finding is that the SPI rate should be as low as possible but still greater than a lower bound which we derive analytically. For phage *λ* we find that the experimentally measured SPI rate matches our model’s prediction. We also derive expressions to describe how stochasticity affects the competition between nonlysogens and lysogens undergoing SPI.

## 2 The LUV model

To understand how SPI plays into the natural selection of lysogenic cells over their uninfected counterparts, we developed a model summarized by equation 1 - 5. This model simulates evolutionary population dynamics using replicator equations [19] with carrying capacity *K* ≈ 10^6^ and in time units of per cell generation (≈ 30 minutes per cell [1]). Since the lysogenic genome contains viral genes, there are many ways in which lysogenized cells could be different from their nonlysogenized counterparts. These differences could influence host cell physiology by directly affecting the intrinsic cell growth rate via expression of viral genes integrated within the host’s genome. Therefore, in our model we assume that the lysogens, *L*, grow at a rate *r* and the nonlysogens, *U*, grow at a rate *g*. The model simulates lysogenic and nonlysogenic cells growing in the same environment with carrying capacity *K* (which reflects nutrient and space limitations). The lysogens undergo SPI at a rate *σ*, and the resulting free viruses, *V*, infect the nonlysogens *U* to induce *α* decision-making events per cell per phage per generation. A proportion *p* of these phage-cell encounters enter the lysogenic pathway, and the remaining proportion 1 − *p* enter the lytic pathway. When the lytic pathway occurs, *b* free viruses are released given by the burst size of the phage. The viruses degrade at rate *γ*.

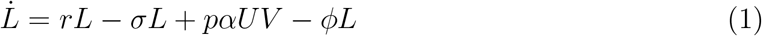

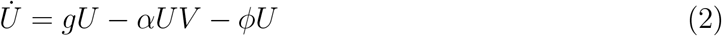

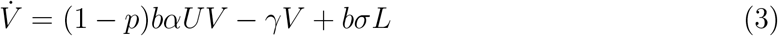

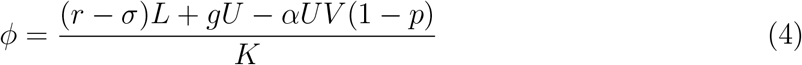

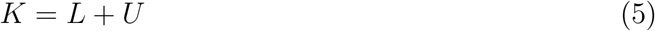

The parameter variables are summarized in detail in Table 1 along with the parameter values for the phage *λ* - *E. coli* system. The burst size of phage *λ* is given by *b* ≈ 150. The probability of lysogeny is known to be host cell volume and MOI dependent, ranging from 20% to 80% [9, 10, 24, 28], but we take *p* ≈ 0.3. Cell growth rates are determined by cell doubling times ranging from 20 mins to 40 mins [1, 29], so we take *g* = 1 cell division per cell per generation. Similarly, we take *r* to be ≈ 1 cell per generation, but we investigate different *r* values in the range 0.9 ≤ *r* ≤ 1 to determine how the growth-cost due to lysogeny affects the dynamics. Phage degradation occurs on the order of 0.1 per day per phage [6], which converts to *γ* ≈ 0.001 per generation per phage. The parameter *α* measures the rate at which nonlysogens are infected and either lysed or lysogenized per generation per phage per cell. This includes the phage infection/adsorption rate and the rate at which infected cells develop lytically or lysogenically per generation. We estimated *α* from the phage adsorption experiment in [4, 17] by fitting a small mass-action model of the phage-cell infection dynamics to the raw data. This *α* estimate allowed fixation events to occur on the timescale of a few days, similar to the experimental results in [7] which compete phage *λ* lysogens against their nonlysogenic counterparts. Increasing *α* by a factor of 10 resulted in fixation events occurring on the timescale of 10s of days, but the overall behavior of the system remained identical. The intrinsic SPI rate of phage *λ* in a *recA^−^* background is < 10^−8^, but in a *recA^+^* background the SPI rate is estimated to be about 10^−6^ to 10^−7^ per generation [11, 12, 29]. Since *recA^−^* mutants probably don’t exist in nature, we use the *recA^+^* SPI rate because natural selection likely acts on the *recA^+^* strain and not the *recA^−^* strain.

**Table 1:** LUV model parameters

| Parameter | Symbol | Value |
| --- | --- | --- |
| Lysogen replication rate | $r$ | $0.9 \leq r \leq 1$ per gen |
| Nonlysogen replication rate | $g$ | 1 per gen |
| Carrying capacity | $K$ | $\geq 10^6$ |
| Phage-cell binding/infection rate | $\alpha$ | $\approx 10^{-7}$ per phage per cell per gen |
| Phage burst size | $b$ | $\approx 150$ |
| Lysogenization rate | $p$ | $\approx 0.30$ |
| Phage degradation rate | $\gamma$ | $\approx 0.001$ per gen |
| SPI rate | $\sigma$ | $\approx 10^{-6}$ per gen |

We can take advantage of the differing time scales of the reactions to impose some conditions (inequalities) on the parameters. From our parameter estimates, we find that the quantity *αbK*/*γ* ≈ 1.5 · 10^4^ which is quite large. The numerator *ν*_1_ = *αbK* is essentially the rate (per generation) at which a population of *K* nonlysogens are lysed or lysogenized given a single phage lytic event released *b* infecting phages. The denominator *ν*_2_ = *γ* is the rate at which a single phage degrades per generation. Since phage degradation is on the order of days and lysis-lysogeny decision-events occur on the order of hours, it is no surprise that the ratio *ν*_1_/*ν*_2_ is quite large using our literature estimates. Thus, we assume that *αbK/γ* ≫ 1 throughout our study which should hold even if there is some uncertainty in the parameter estimates since this quantity is so large compared to 1. We also assume that *r* > *σ* because if *σ* > *r* the net growth rate of the lysogens would be negative indicating an unsustainable population. The lysogen’s genome is the same as the nonlysogen’s genome except that it contains more genes from the integrated viral genome (the genes from the prophage). Maintenance of the lysogenic state generally requires continued gene expression of a key viral repressor protein to block entrance into the lytic pathway, and in the specific case of phage *λ* this protein is the CI repressor. It is possible that the cell incurs some growth-rate cost by maintaining expression of this viral repressor. Experiments competing *λ* lysogens against their nonlysogenic counterparts showed that their growth rates are not statistically different, suggesting that *r* = *g* [7]. However, it is still possible that *r* could be different from *g* by a very small amount that could be difficult to detect. For example, this could happen if *g* − *r* = 0.01 or less. From these observations we assume that the nonlysogenic growth rate *g* is at least *r* or otherwise at least slightly larger. These inequalities will be important later on when analyzing this model. Note that since *g* ≥ *r* then *g* > *σ* as well. Also note that if *r* > *g* then the lysogens would displace the nonlysogens even without SPI in the long run, and the inclusion of SPI when *r* > *g* would only accelerate the process. Since this observation is somewhat trivial, we do not specifically investigate it in this work. However, we do prove this analytically later on.

With the model and its parameters specified, we numerically simulated the LUV model using the Runge Kutta integration scheme (see Fig. 1) for the parameter values in Table 1 to gain an understanding of the role of SPI in the competition between lysogens and nonlysogens. The initial conditions we chose were *L*(*t* = 0) = 0.01*K* and *U* = 0.99*K* which implies that the lysogens initially comprise 1% of the population. Since *K* ≥ 10^6^ the initial values of *L* and *U* were at least 10^4^ which allows us to well-approximate the behavior of this system using deterministic models. We investigate stochastic models later on in this work given the fact that the low SPI rate can still introduce stochasticity into the model. The simulations show that this system tends towards a state in which the lysogens completely displace the nonlysogens in the long run even though the lysogenic growth rate is lower (*g* > *r*). This suggests that SPI allows the lysogens to subvert the nonlysogens even if the lysogenic state is costly to cell growth. This result is consistent with experiment observations in which lysogen strains displaced lysogen-cured strains when growth together [3, 7, 18]. Essentially, a low but steady rate of lysogens undergoing SPI allows the released phages to either kill off the nonlysogen strain or convert them to the lysogenic state.

**Figure 1:**
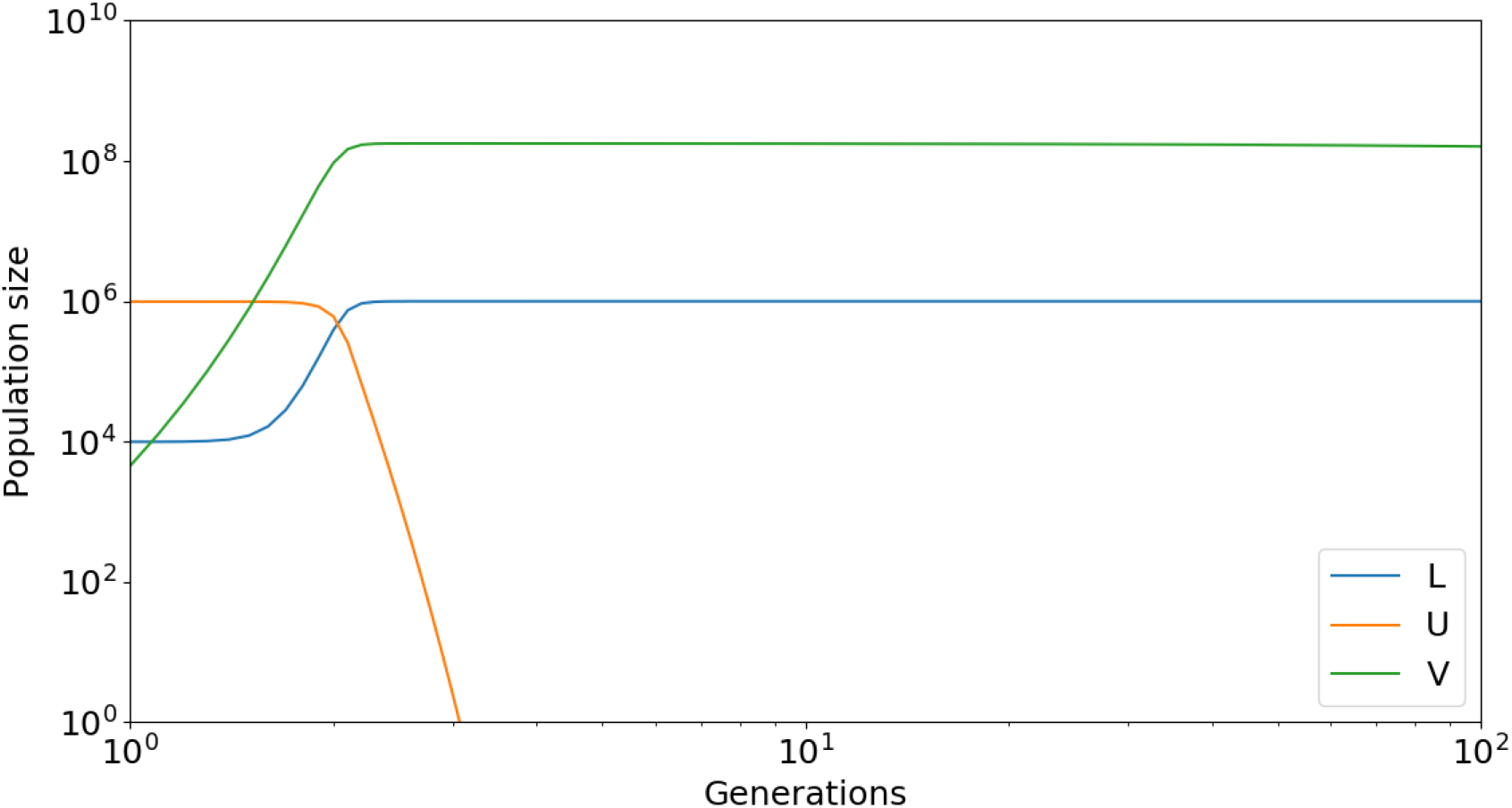
LUV model simulated using parameters from Table 1. Specifically, we set *r* = 0.99 and *σ* = 10^−6^. We get the same results if we vary the model’s parameters.

## 3 Lower bound on SPI rate for lysogenic advantage

To determine the exact bounds on the SPI rate necessary for the lysogens to displace the nonlysogens, we analytically solved our model (equations 1 - 5) at steady state to obtain its equilibrium points and then tested their stability by calculating the eigenvalues of the system’s associated Jacobian matrix. At first glance, this seems like a very difficult task due to the nonlinearity of the model, but fortunately the model admits analytical solutions for its equilibrium points. We used Mathematica to check all calculations and derivations.

First, we set 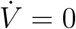 in equation 3 and solve for *V*. This gives us the following relationship.

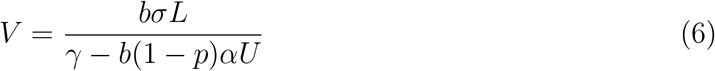

Next, we set 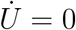 using equation 2.

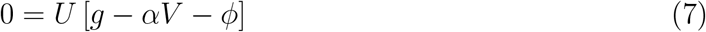

The only way for 7 to be zero is if either *U* = 0 or if [*g* − *αV* − *ϕ*] = 0. We will analyze both cases.

If *U* = 0 it immediately requires that *L* = *K* by constraint equation 5. This then immediately implies that *V* = *bσK*/*γ* by equation 6. Thus, the point 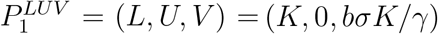 is one equilibrium point of our system.

Otherwise, if *g* − *αV* − *ϕ* = 0 then we can plug in 4 for *ϕ*. The relation *g* − *αV* − *ϕ* = 0 immediately implies that equation 8 must hold.

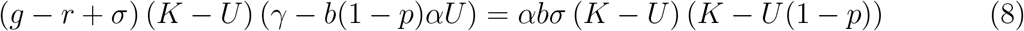

We can immediately note that *U* = *K* is a possible solution of equation 8. Setting *U* = *K* implies *L* = 0 by the constraint 5. Then, using equation 6 we immediately see that *V* = 0 if *U* = *K*. Thus, the second equilibrium point of our system is given by 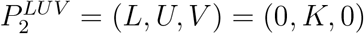.

If *U* ≠ *K*, then we can cancel the factor (*K* − *U*) and solve for *U* to obtain *U* = *γ* (*g* − *r* − *σ* (*αbK/γ* − 1))/*αb*(1 − *p*)(*g* − *r*). We can then use this value for *U* to calculate *L* and *V* using *L* = *K* − *U* and equation 6. This solution set defines equilibrium point 3, which we denote using 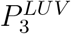.

The equilibrium points of our system are summarized in Table 2. To summarize, 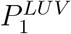 describes a state in which the lysogens have displaced the nonlysogens, 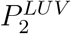 describes a state in which the nonlysogens have displaced the lysogens, and 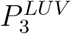 describes a state in which both the lysogens and nonlysogens coexist. Given these equilibrium points, we seek to determine the conditions necessary for the lysogens to displace the nonlysogens.

**Table 2:** LUV model equilibrium points

| Equilibrium point | L | U | V |
| --- | --- | --- | --- |
| $P_1^{LUV}$ | $K$ | $0$ | $b\sigma K/\gamma$ |
| $P_2^{LUV}$ | $0$ | $K$ | $0$ |
| $P_3^{LUV}$ | $K - \frac{g-r-\sigma(\alpha b K/\gamma-1)}{(1-p)(g-r)\alpha b/\gamma}$ | $\frac{g-r-\sigma(\alpha b K/\gamma-1)}{(1-p)(g-r)\alpha b/\gamma}$ | $\frac{(g-r+\sigma)-\alpha b K((g+r)(1-p)-\sigma)/\gamma}{\alpha(1+\alpha b K)(1-p)/\gamma}$ |

### 3.1 Stability of equilibrium points for LUV model

To determine the conditions required for the lysogens to displace the nonlysogens, we investigated the stability of the LUV model’s equilibrium points (Table 2) by computing the eigenvalues of our model’s associated Jacobian matrix *J^LUV^*.

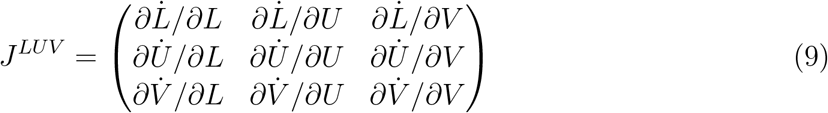

We calculate the partial derivatives in *J^LUV^* using equations defining the LUV model (equations 1 - 5) and we define 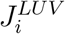 to be *J^LUV^* evaluated at equilibrium point 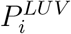.

First, we analyze the stability of equilibrium point 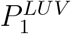 from Table 2, namely (*L, U, V*) = (*K*, 0, *bσK/γ*). This point characterizes a state of the system in which the lysogens have displaced the nonlysogens. Using these values for *L, U*, and *V* we calculate 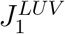.

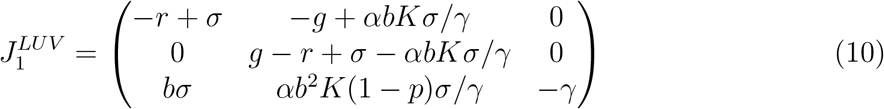

The eigenvalues of 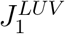 are given by calculating the determinant 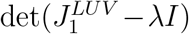 where *I* is the identity matrix. This results in the following characterisic polynomial with eigenvalue *λ* (equation 11).

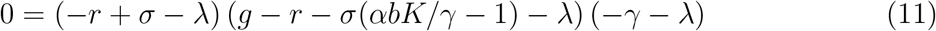

The eigenvalues are the values of *λ* which solve equation 11. We see that these values are *λ*_1_ = −*r* + *σ, λ*_2_ = *g* − *r* − *σ*(*αbK/γ* − 1), and *λ*_3_ = −*γ*. The equilibrium point (*K*, 0, *bKσ/γ*) is a stable point of the system if all eigenvalues are negative. Since *γ* > 0 we know that *λ*_3_ is negative. Since *σ* < *r* we have that *λ*_1_ is negative since *λ*_1_ = −(*r* − *σ*) < 0. The eigenvalue *λ*_2_ is negative if and only if *σ* > *σ_LB_* with *σ_LB_* given by equation 12.

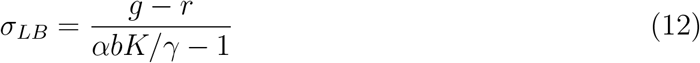

Using literature estimates for the parameter values from Table 1, and specifically with *r* = 0.99 and *σ* = 10^−6^, we estimate *σ_LB_* ≈ 6 · 10^−7^. Thus, *σ* must be larger than this lower bound (defined as *σ_LB_*) in order to establish (*K*, 0, *bKσ/γ*) as a stable equilibrium point. This is very close to the experimentally measured value between 10^−6^ or 10^−7^ per generation for phage *λ* measured most recently [29].

Recall that we assume *g* ≥ *r* so there are two possibilities, either *g* = *r* or *g* > *r*. If *g* = *r* then the lysogenic state does not hurt cell growth rate and, as a consequence, any *σ* > *σ_LB_* = 0 would cause the point (*K*, 0, *bKσ/γ*) to be a steady equilibrium point. Furthermore, if *g* = *r* then critical point 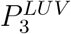 no longer exists (the system goes from having 3 equilibrium points to only 2). Otherwise, if *g* > *r* then the factor *g* − *r* is greater than 0 and hence makes the lower bound *σ_LB_* nonzero. Using the values in Table 1 we estimated that if *g* − *r* ≈ 0.01 per generation then *σ_LB_* ≈ 6 · 10^−7^ per generation. If the lysogenic growth-cost was even smaller (e.g. if *g* − *r* is smaller), then *σ_LB_* < 10^−7^. For example, with *g* = 0.999 then *σ_LB_* = 6 · 10^−8^. The literature estimate for *σ*\* is 10^−6^ [29] which is ≥ *σ_LB_* even in the case that *r* = 0.99. This literature estimate is roughly 10 times larger than this lower bound, which means *σ*\* > *σ_LB_*, implying that equilibrium point 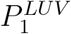 is a stable state and equilibrium point 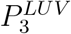 does not exist. Note that if *g* − *r* was smaller (e.g. the lysogenic and nonlysogenic growth rates were very similar, which is what we expect) or if *K* was larger (say *K* = 10^9^), this lower bound would be even smaller causing *σ_LB_* to tend towards 0. Thus, the literature estimate of the phage *λ* SPI rate *σ*\* is likely much larger than the lower bound *σ_LB_*. The important thing to note here is that this lower bound is easily a very small number due to the size of the constants in the denominator. Although we assume *g* ≥ *r*, for completeness we also examine what happens when *r* > *g*. In this case the lower bound *σ_LB_* is negative. Thus, if *r* > *g* then the lysogens will displace the nonlysogens even if *σ* = 0 because 0 > *σ_LB_*.

Next, we examine equilibrium point 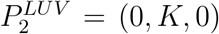. This point describes a state in which the nonlysogens have displaced the lysogens. The stability of point 2 is assessed by calculating the eigenvalues of 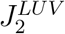 matrix. The Jacobian matrix for (*L, U, V*) = (0, *K*, 0) is given in equation 13.

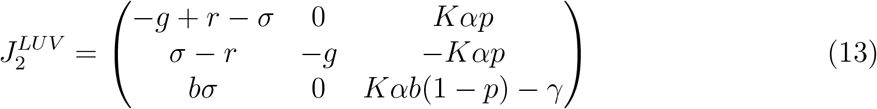

The eigenvalues of 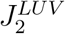 are obtained by calculating its determinant, which results in a characteristic polynomial (equation 14).

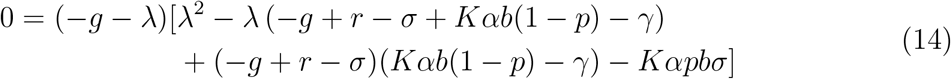

The eigenvalues are then obtained in a straightforward manner. First, *λ*_1_ = −*g* < 0. For *λ*_2_ and *λ*_3_, we see that these are the roots of the quadratic inside the square brackets in equation 14. The roots of this quadratic take the form 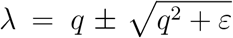 with *q* = (−*g* −*γ* + *αbK*(1 − *p*) + *r* − *σ*)/2 and *ε* = ((*αbK*(1 − *p*) − *γ*)(*g* − *r*) + (*αbK* − *γ*)*σ*)). It is easy to show that if *ε* > 0 then one of the solutions *λ* > 0. In this case, we have *ε* > 0 because every factor and term in the *ε* expression is positive. We have that (*αbK/γ* − 1) > 0, (*g* − *r*) ≥ 0, *γ* > 0, and *σ* > 0. Since *αbK/γ* ≫ 1 we also have that (1 − *p*)*αbK/γ* > 1. Thus, equilibrium point 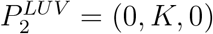 is an unstable equilibrium point of this system.

Finally, we examine the stability of 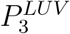 which describes a state of the system in which both the lysogens and nonlysogens coexist. First, note that this equilibrium point does not exist if *σ* is larger than the lower bound *σ_LB_* in 12. Thus, if inequality 12 holds then we do not need to analyze point 3’s stability. Since the literature estimate *σ*\* > *σ_LB_* we hypothesize that this equilibrium point indeed does not exist, but for completeness we examine its stability here.

If *σ* < *σ_LB_* then both 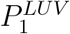 and 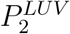 are unstable. If a steady state indeed does exist under the condition *σ* < *σ_LB_* then it must be true that 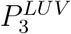 becomes the (only) steady state of the system. The Jacobian matrix for this equilibrium point, 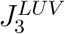, is a 3 by 3 matrix with extremely complicated expressions for its elements due to the fact that the expressions for *L, U*, and *V* for 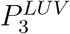 are quite complicated themselves. Therefore, we do not show the full Jacobian here, but we numerically evaluate its eigenvalues by using parameter values from Table 1 except we set *σ* = *θ* · *σ_LB_* so that inequality 12 does not hold, which means *θ* < 1. We discovered that for all *θ* < 1 the eigenvalues of 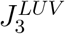 all had strictly negative real parts, implying that this becomes a steady state of the system if *σ* < *σ_LB_*. Therefore, if *σ* is extremely small (so that the SPI rate is lower than *σLB*) then the steady state of the system is one in which both the lysogens and nonlysogens coexist.

Overall, we have shown that lysogens will displace nonlysogens even if the lysogenic state is costly to cell growth provided that *σ* is at least (*g* − *r*)/[*Kαb/γ* − 1]. We have also shown that the true SPI rate of phage *λ* is likely greater than this lower bound, suggesting that the SPI rate has evolved so that lysogens can displace nonlysogens. However, this analysis only tells us the lower bound on *σ* required for lysogens to have the advantage. That is, *σ* is allowed to be any number greater than *σ_LB_*. Thus, it is natural to ask if a larger SPI rate is advantageous.

## 4 The LUV2 model

In the previous section we showed that if *r* > *σ* > (*g* − *r*)/ [*Kαb/γ* − 1] = *σ_LB_* then a mixture of lysogenic and nonlysogenic cells will tend towards a state in which the nonlysogens are displaced in favor of the lysogens. Note that this inequality places no strong restriction on the exact value of *σ*. The only requirement is that it must be strictly greater than *σ_LB_* and must be less than the intrinsic cell replication rate *r*. To understand what value of *σ* may be optimal, we adjusted the LUV model so that it included two lysogen strains, *L*_1_ and *L*_2_, which differ only in their SPI rates *σ*_1_ and *σ*_2_ respectively. These lysogens will compete with each other and with the nonlysogenic cells *U*. These lysogen strains also produce their own phages, denoted *V*_1_ and *V*_2_, respectively. All other parameters and interactions between the two lysogen strains and the nonlysogens is kept the same so that we ensure we are only comparing different SPI rates. The overall idea here is that we want to identify an SPI rate which would allow a lysogen strain to outcompete another lysogen strain while also being able to displace the nonlysogens. The new model is summarized by equations 15 - 21.

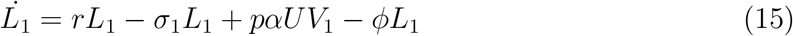

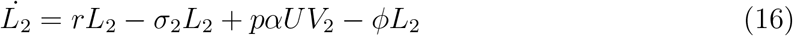

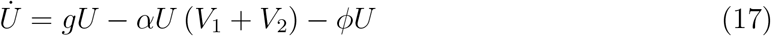

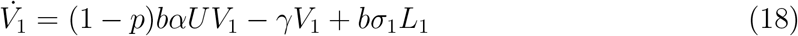

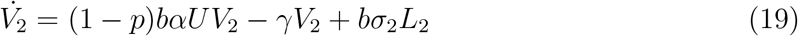

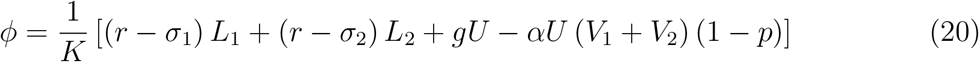

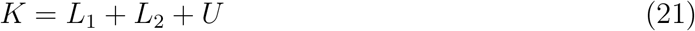

In this model, *L*_1_ denotes the number of lysogen strain 1 with SPI rate *σ*_1_ and *L*_2_ denotes the number of lysogen strain 2 with SPI rate *σ*_2_. The variables *V*_1_ and *V*_2_ are the phages emitted from lysogen strains 1 and 2, respectively, via the SPI process. All parameters are the same as in the previous model. Without loss of generality, we assume that *σ*_1_ > *σ*_2_ so that lysogen strain 1 has a higher SPI rate than lysogen strain 2. Note that here we are also assuming the same inequalities hold from analyzing the LUV model, namely *αbK/γ* > 1, *g* − *r* ≥ 0, and *r* > *σ*_1_. Since *σ*_1_ > *σ*_2_ we also have *r* > *σ*_2_.

To gain an understanding of the behavior of this system, we simulated it using the parameter values in Table 1. We *r* = 0. 99 and *σ*_1_ to be 10 times greater than *σ*_2_ with *σ*_2_ = 10^−6^. In Fig 2 we show a sample trajectory of the system using initial conditions *L*_1_(*t* = 0) = 0.005*K, L*_2_(*t* = 0) = 0.005*K*, and *U*(*t* = 0) = 0.99*K* so that the lysogens initially only comprise 1% of the population, and this 1% is split evenly among the two lysogen strains. We see that early on, *L*_1_ has a clear advantage in the sense that it greatly outnumbers both *L*_2_ and the nonlysogens *U*. However, in the long run we see that *L*_2_ actually starts to overtake *L*_1_, and that this switch occurs after *U* decreases towards 0.

**Figure 2:**
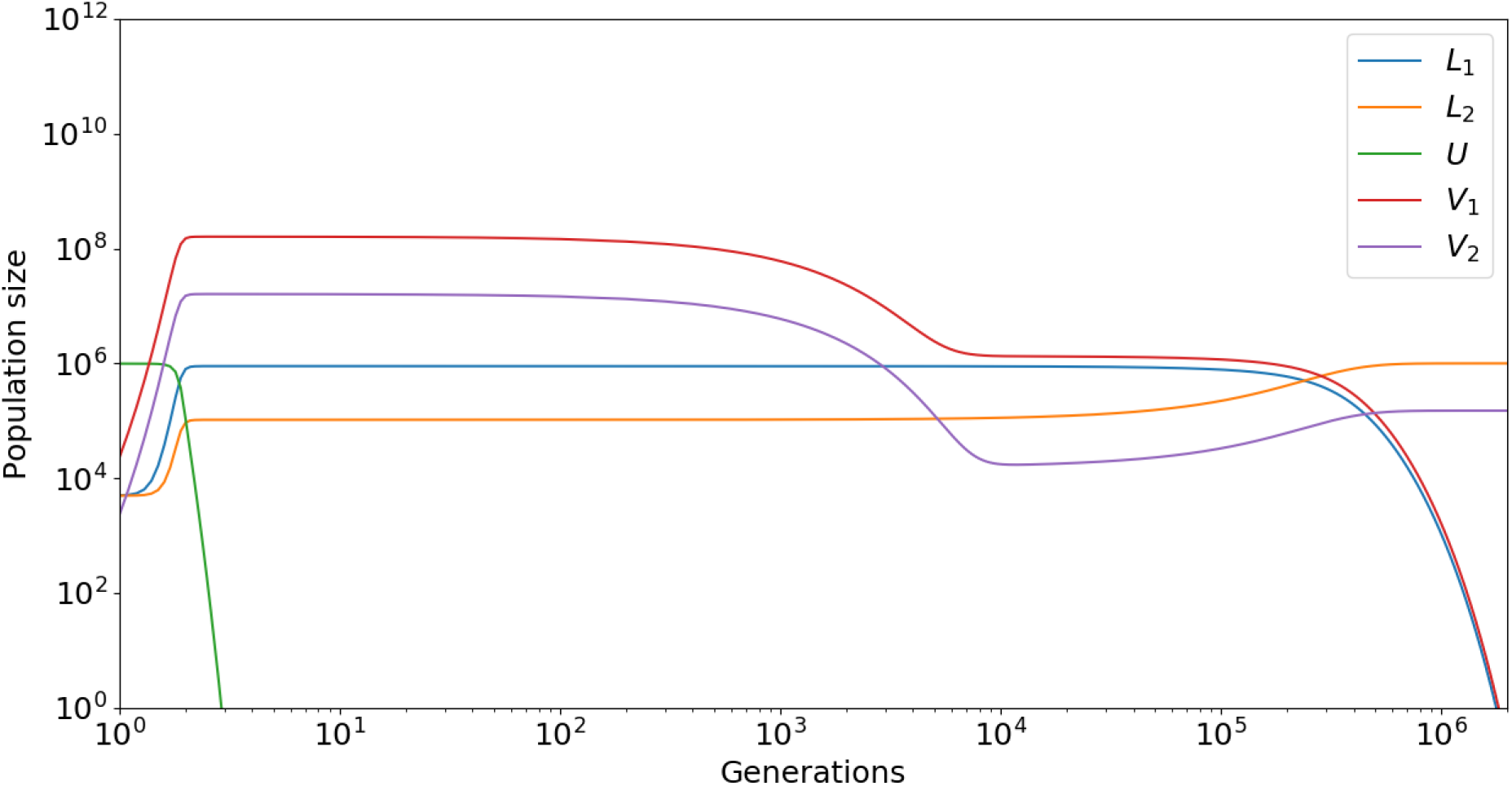
LUV2 model simulation using default parameters from Table 1. We used *r* = 0.99, *σ*_2_ = 10^−6^, and *σ*_1_ = 10^−5^.

Numerical simulations of various parameter sets showed similar behavior, with *L*_2_ always displacing both *L*_1_ and *U* in the long run. This suggests that lower SPI rates are favorable in the long run. Intuitively, we can reason that once the nonlysogens go extinct there is no more benefit to having a higher SPI rate. Thus, we can conclude that if both lysogen strains *L*_1_ and *L*_2_ survive after *U* reaches 0, then the lysogen with the lower SPI rate will be selected for in the long run.

## 5 Slower SPI rates are advantageous

To prove that the lysogenic strain with the lower SPI rate is the one which is favored by natural selection, we calculate the equilibrium points of our model and assess their stability. We impose the requirement that each of these lysogen strains be able to outcompete nonlysogens if separately competed with them, which implies that both *σ*_1_ > *σ_LB_* and *σ*_2_ > *σ_LB_*. As in the LUV model, we first begin with setting the free phage rate equations to zero (equations 18 and 19) i.e. 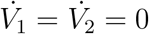. This results in the following relationships for *V*_1_ and *V*_2_.

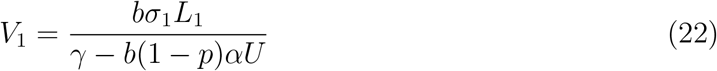

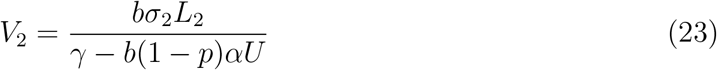

Next, we set equation 17 to 0 so that 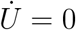. This implies that:

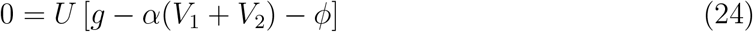

Equation 24 can be solved if either *U* = 0 or if *g*−*α*(*V*_1_+*V*_2_)−*ϕ* = 0. We will examine both cases. If *U* = 0 then we immediately see that *V*_1_ and *V*_2_ are determined by their respective lysogen strain levels. That is, if *U* = 0 we have that *V*_1_ = *bσ*_1_*L*_1_/*γ* and *V*_2_ = *bσ*_2_*L*_2_/*γ*. Thus, we focus on *L*_1_ and *L*_2_ next. If *U* = 0 then *K* = *L*_1_ + *L*_2_ by the constraint 21. This means that *L*_1_ = *K* − *L*_2_. We can plug this into the rate equation for 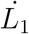 (equation 15) and set 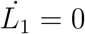. After simplifying, this results in an equation with only 1 unknown variable, namely *L*_2_.

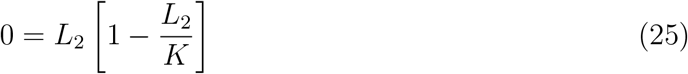

The two solutions to this equation are *L*_2_ = 0 or *L*_2_ = *K*. Along with the constraint (equation 21), we know that (*L*_1_, *L*_2_) = (*K*, 0) and (*L*_1_, *L*_2_) = (0, *K*) are both solutions of equation 25. Using equations 6 and 23, we can write down two equilibrium points of our system (*L*_1_, *L*_2_, *U, V*_1_, *V*_2_), namely 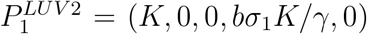 and 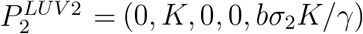.

If instead *g* − *α*(*V*_1_ + *V*_2_) − *ϕ* = 0 then we can combine this equation with equations 22, 23, and 20 to show that the following equation must hold.

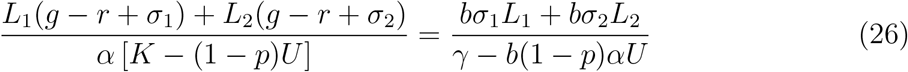

We can immediately see that if *L*_1_ = *L*_2_ = 0 then equation 26 holds. If *L*_1_ = *L*_2_ = 0 then *U* = *K* by the constraint 21. Furthermore, if *L*_1_ and *L*_2_ are both zero then we also have that *V*_1_ = *V*_2_ = 0 by equations 6 and 23. Thus, the third equilibrium point of the system is given by 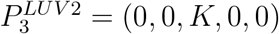.

If both *L*_1_ and *L*_2_ are not zero, then we need to solve the equation 26 for both *L*_1_, *L*_2_, and *U* along with equations 15 - 21. However, this is very difficult to do by hand. Thus, we use Mathematica here to derive the additional solutions. It turns out that there are three additional solutions, which we denote using 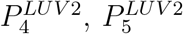, and 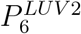. However, some of these solutions are extremely complicated expressions, so we do not show them in their full detail. These additional solutions describe states of the system in which *L*_1_, *L*_2_, and *U* coexist in the long run. Hence, these solutions are not of particular interest since our goal is to determine the conditions required for a lysogen strain to displace all others and the nonlysogens.

For equilibrium points 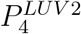 and 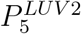, we find that the expression for *U* has the same form, namely *U* = [*γ*(*g* − *r* + *σ_j_*) − *αbKσ_j_*)] /(*αb*(1 − *p*)(*g* − *r*)) with *j* = 1 for point 4 and *j* = 2 for point 5. In order for these equilibrium points to be unique (i.e. different from points 1,2 and 3) we must have that 0 < *U* < *K*. We find that *U* > 0 if and only if *σ_j_* < (*g* − *r*)/ [*Kαb/γ* − 1] = *σ_LB_*. However, this contradicts our initial assumption that both *σ*_1_, *σ*_2_ > *σ_LB_*. Indeed, *σ*_1_ > *σ_LB_* and *σ*_2_ > *σ_LB_* implies that *U* < 0 which is unphysical since population counts cannot be negative. Therefore, equilibrium points 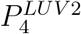 and 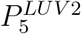 do not exist under our parameter regime.

Finally, we analyze the equilibrium point 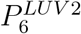 for which expressions for *L*_1_, *L*_2_, and *U* are as follows:

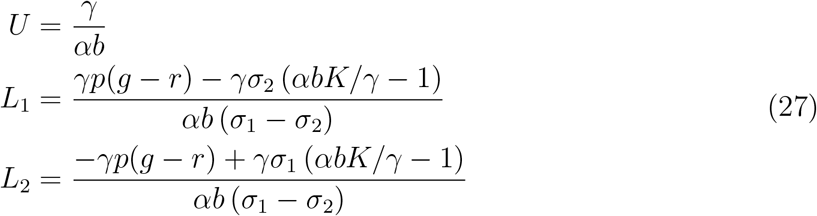

Using the fact that (*σ*_1_ − *σ*_2_) > 0 we can easily show that *L*_1_, *L*_2_ > 0 if and only if *σ*_1_ > *p*(*g* − *r*)/ [*Kαb/γ* − 1] and *σ*_2_ < *p*(*g* − *r*)/ [*Kαb/γ* − 1]. These conditions are consistent with *σ*_1_ > *σ*_2_, but they contradict the fact that *σ*_2_ > *σ_LB_*. To see this, note that *σ*_2_ < *p*(*g* − *r*)/ [*Kαb/γ* − 1] < (*g* − *r*)/ [*Kαb/γ* − 1] = *σ_LB_* since 0 < *p* < 1. Indeed, *σ*_2_ > *σ_LB_* implies that *L*_1_ < 0 which is unphysical. Therefore, our original requirement that *σ*_1_, *σ*_2_ > *σ_LB_* causes equilibrium point 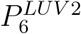 to not exist.

All equilibrium points of the LUV2 model which exist under our parameter regime are summarized in Table 3. Note that these equilibrium points are similar to the ones we found for the LUV model.

**Table 3:**
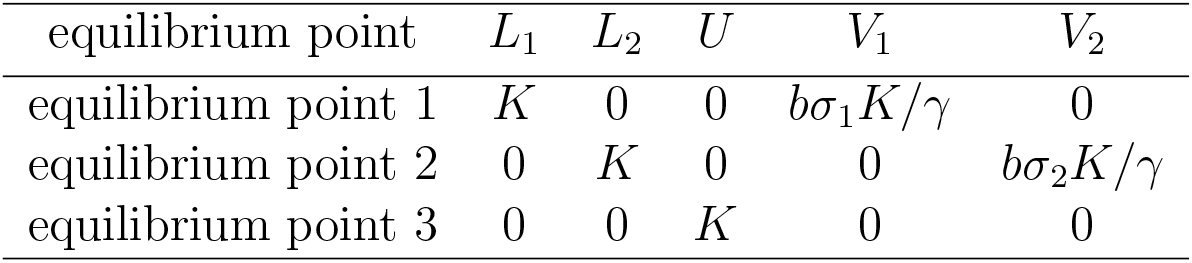
LUV2 model equilibrium points

### 5.1 Stability analysis of LUV2 equilibrium points

To understand what conditions enable one lysogen strain to outcompete the other, we analyzed the stability of our model’s equilibrium points by calculating the eigenvalues of the Jacobian matrix *J^LUV2^* for each equilibrium point in Table 3.

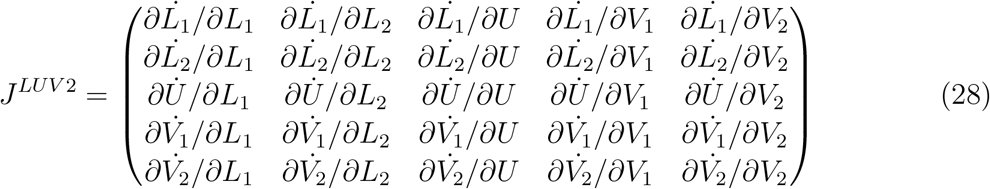

For 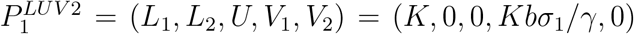, the Jacobian matrix is given by 29. This equilibrium point describes a state in which lysogen strain *L*_1_ (with the higher SPI rate) displaces both the nonlysogens *U* and lysogen strain *L*_2_.

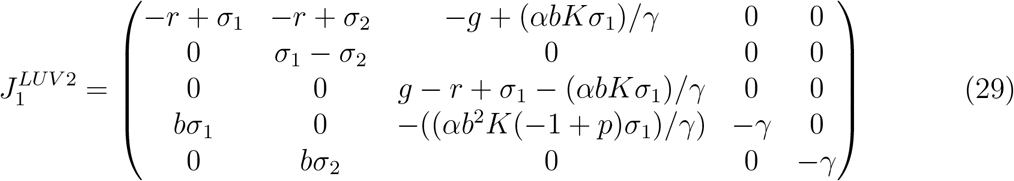

The eigenvalues of 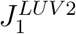 are determined by solving the equation 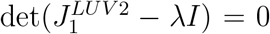 which results in the following characteristic polynomial.

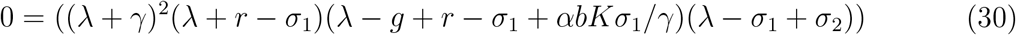

The eigenvalues are then easily obtained by setting each factor to 0 and solving for *λ*. The eigenvalues are *λ*_1_ = −*γ, λ*_2_ = −*γ, λ*_3_ = −*r* + *σ*_1_, *λ*_4_ = *g* − *r* − *σ*_1_(*αbK/γ* − 1), and *λ*_5_ = *σ*_1_ − *σ*_2_. Without inspecting each eigenvalue, we can immediately conclude 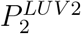 is an unstable equilibrium point because eigenvalue *λ*_5_ > 0 since *σ*_1_ > *σ*_2_. Thus, the system will not tend to a state in which lysogen strain *L*_1_ displaces lysogen strain *L*_2_ and the nonlysogens *U*.

Next, we analyze equilibrium point 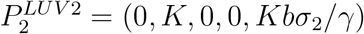. This point describes a state in which lysogen strain *L*_2_ (with the lower SPI rate) displaces both the nonlysogens *U* and lysogen strain *L*_1_. The corresponding Jacobian is given by 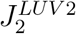 in equation 31.

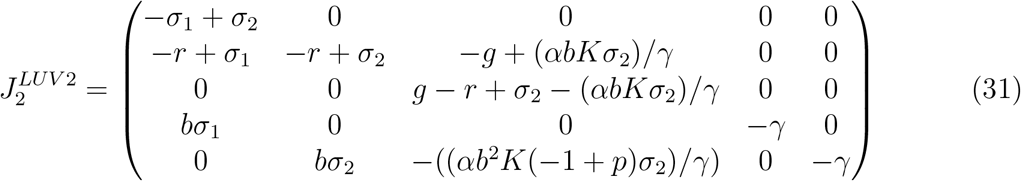

The corresponding characteristic equation is given by:

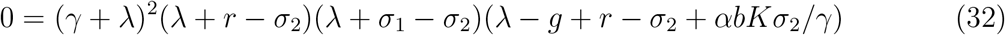

The eigenvalues are given by *λ*_1_, *λ*_2_ = −*γ, λ*_3_ = −*r* + *σ*_2_, *λ*_4_ = −*σ*_1_ + *σ*_2_, and *λ*_5_ = (*g* − *r*) − *σ*_2_ (*αbK/γ* − 1). Eigenvalues 1 - 4 are obviously negative since *γ* > 0, *σ*_1_ > *σ*_2_, and *σ*_1_, *σ*_2_ < *r*. The inequality *σ*_2_ > *σ_LB_* can be rewritten as 0 > (*g* − *r*) − *σ*_2_ (*αbK/γ* − 1) which shows that 0 > *λ*_5_. Therefore, all eigenvalues are negative and therefore equilibrium point 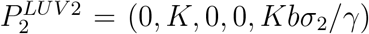 is a stable equilibrium point of the system. Thus, this system will tend towards a state in which lysogen strain *L*_2_ displaces lysogen strain *L*_1_ and the nonlysogens *U*.

Finally, we examine equilibrium point 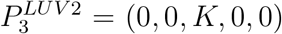 which describes a state in which both lysogen strain *L*_1_ and *L*_2_ are displaced by the nonlysogens *U*. The Jacobian for this equilibrium point is given by 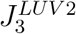.

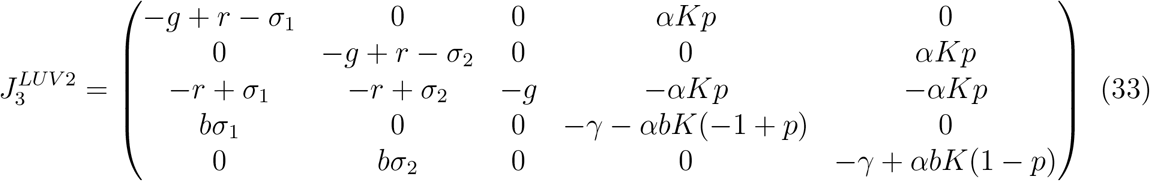

The characteristic equation is given by:

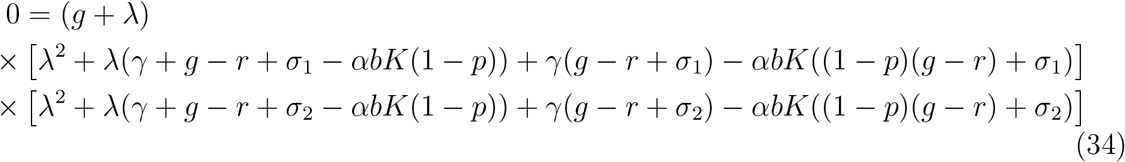

The first eigenvalue is given by (*g*+*λ*) = 0 ⇒ *λ*_1_ = −*g* < 0. The remaining eigenvalues are given by solving the quadratic equations within both square brackets [·]. These eigenvalues are 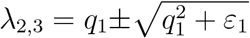 and 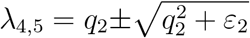, with *q_j_* = (−*g* − *γ* + *αbK*(1 − *p*) + *r* − *σ_j_*)/2 and *ε_j_* = −*gγ* + *αbK*((*g* − *r*)(1 − *p*) + *σ_j_*) + *γr* − *γσ_j_* for *j* = 1, 2. If *ε_j_* > 0 then it must be true that at least one of the eigenvalues is positive, implying that this equilibrium point is not a stable steady state. It is straightforward to show that *ε_j_* < 0 only if we have *σ_j_* < − (*g* − *r*) [*αbK*(1 − *p*)/*γ* − 1] / [*αbK/γ* − 1] < 0. However, we require *σ_j_* > *σ_LB_* > 0 ⇒ *ε_j_* > 0 and hence this equilibrium point is unstable. Thus, the system will not tend towards a state in which the nonlysogens dipsplace both lysogen strains *L*_1_ and *L*_2_.

Overall these results collectively show that *L*_2_ will displace both *L*_1_ and *U* if *σ_LB_* < *σ*_2_ < *σ*_1_. The upper half of the inequality *σ*_2_ < *σ*_1_ gives *L*_2_ a competitive advantage against *L*_1_, and the lower half of the inequality *σ_LB_* < *σ*_2_ gives *L*_2_ the competitive advantage against the nonlysogens. Thus, a lower SPI rate is advantageous over a higher SPI rate in the long run.

## 6 Stochastic factors influencing selection of SPI rate

To understand how stochastic factors could influence the competition between lysogens and nonlysogens, we simulated the LUV model with some modifications. Since the SPI rate is fairly small, at low lysogen counts (e.g. *L* ≈ 100 or less) it is possible for several generations to pass before a single SPI reaction takes place. This is difficult to capture in the LUV model because it is simulated using ODEs which are deterministic. In this deterministic setting, some SPI reactions occur from the start of the dynamics since the net rate of SPI, *L* · *σ*, is > 0 at *t* = 0. Thus, we modified our LUV model so that the SPI reactions would occur stochastically during the dynamics. Over an small time interval Δ*t* over which we integrate, the probability that a single lysogen undergoes SPI is *σ* · Δ*t*. For *L* lysogens, the probability that *ℓ* < *L* of them undergo SPI in Δ*t* is given by a binomial distribution as in equation 35 with the average number of SPI events in a time interval Δ*t* given by 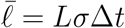.

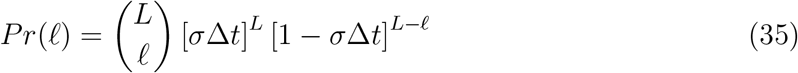

In Figure 3 we show 2 sample simulations of our model at an initial lysogen count of *L*_0_ = *L*(0) = 0.001 · *K*. In the right panel we observe the lysogens being slowly displaced by the nonlysogens (since *r* < *g*) but when an SPI reaction occurs this “saves” the lysogens and they end up taking over the population. However, it can also happen that the SPI reaction does not occur at all. If an SPI reaction fails to fire early enough (as in the left panel), the probability that an SPI reaction will fire later approaches 0. This is because the probability of an SPI reaction is proportional to *L* · *σ*, and since *σ* is already a very low number (≈ 10^−6^) having low *L* makes this probability approach 0. Thus, if stochasticity plays a major role (e.g. at low population counts) and if *r* < *g*, then the total SPI rate, *L* · *σ*, must not be too low if the lysogens are to displace the nonlysogens.

**Figure 3:**
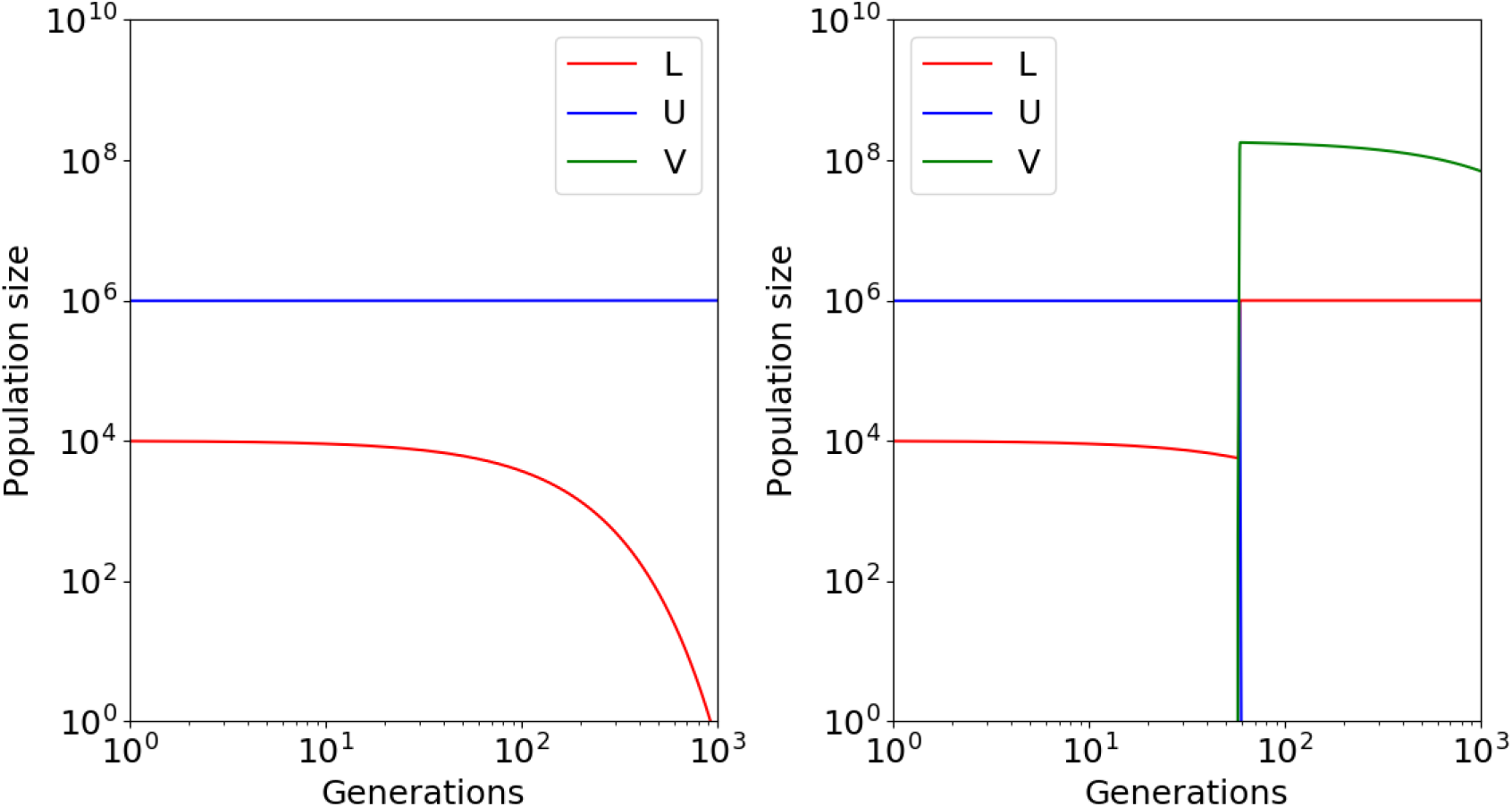
Stochastic LUV model simulations using the parameters from Table 1. In the left panel, the nonlysogens displaced the lysogens because no SPI reaction occurred. In the right panel an SPI reaction occurred early enough for the lysogens to displace the nonlysogens. We used parameter values from Table 1 with *L*(0) = 0.001 · *K* = 1000. We also set *r* = 0.99, *σ*_1_ = 10^−5^, and *σ*_2_ = 10^−6^.

At high initial lysogen counts, the lysogens almost always displaced the nonlysogens because an SPI event almost always occurs. On the other hand, when the initial lysogen count is low (e.g. 10 or 100) then they often were displaced by the nonlysogens. A similar trend was observed for *σ*, namely that high *σ* allowed the lysogens to displace the nonlysogens more frequently relative to the case of a very low *σ* value. Additionally, if the lysogenic growth-cost was small (e.g. *g* ≈ *r*), then the lysogens were able to displace the nonlysogens with greater probability over a much wider range of L(0) and *σ* values. To systematically understand the effect of *L*(0), *σ*, and *r* on the probability that the lysogens displace the nonlysogens, we estimated the probability of lysogen-sweeping events from repeated runs of the stochastic simulation. In Figure 4 we show the probability that *L* = *K, U* = 0 over a range of *L*(0) and *σ* values for *r* = 0.9 and *r* = 0.999. Generally we see that at low *L*(0) and *σ* values the lysogens usually cannot displace the nonlysogens. On the other hand, at higher *L*(0) and *σ* the lysogens were always able to more probably displace the nonlysogens. For intermediate values of *L*(0) and *σ*, the probability that the lysogens displace the nonlysogens ranges from 0 to 1. But as *r* approaches *g* (right panel with *r* = 0.999), we see that the lysogens can displace the nonlysogens over a much wider range of *L*(0) and *σ* values, even if *L*(0) and *σ* are low. When *g* − *r* = 0, the lysogens always displace the nonlysogens even in stochastic simulations. This is because the two populations remain at their initial values until the first SPI reaction occurs, no matter how long it takes. Thus, if the lysogenic growth-cost is small or zero, we expect that the lysogens will generally displace the nonlysogens with high probability.

**Figure 4:**
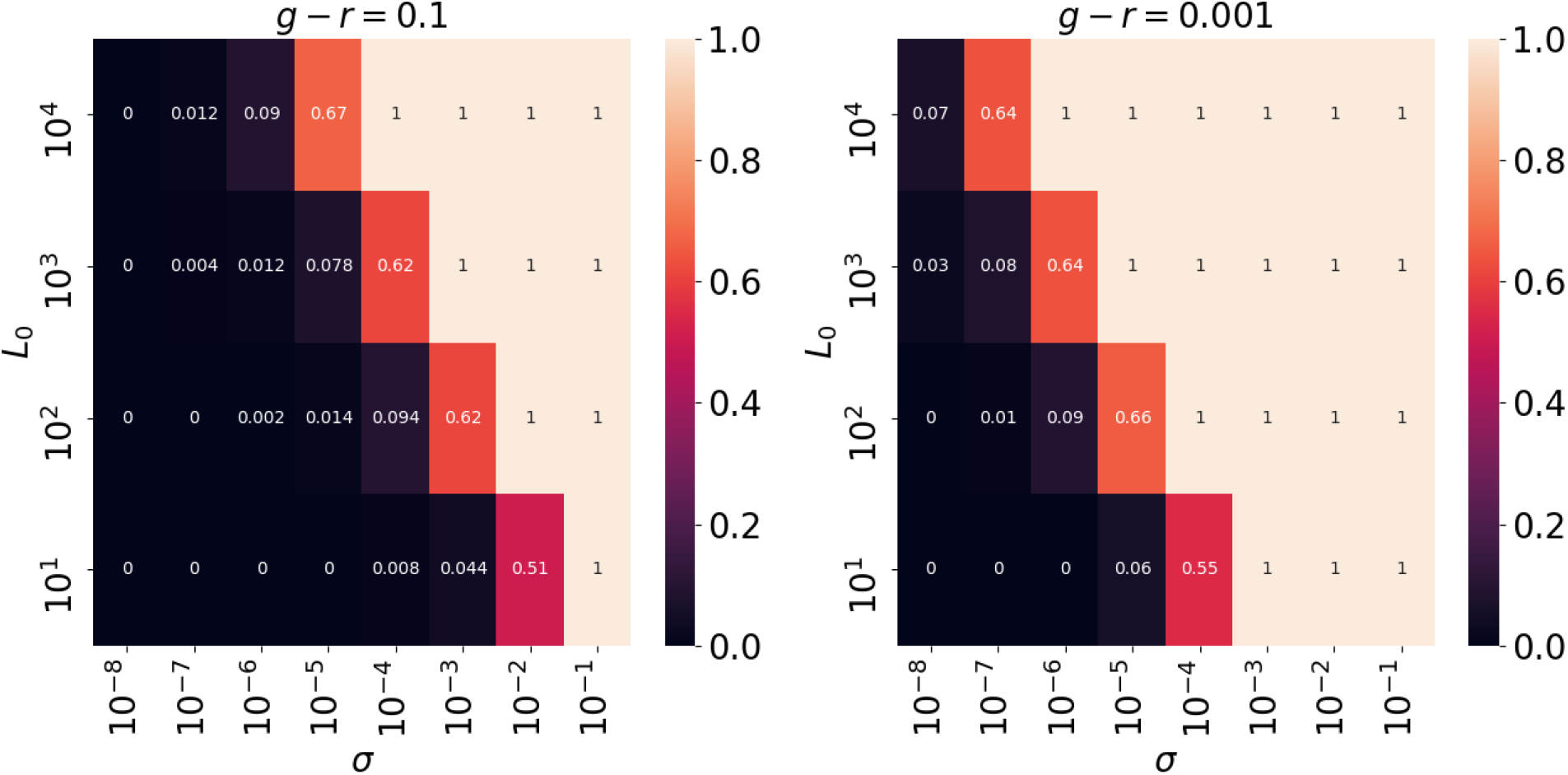
We calculated the fraction of simulations resulting in lysogens displacing the nonlysogens for different values of *L*(0) and *σ* using the stochastic LUV model. The left panel was simulated at *r* = 0.9 and the right panel was simulated at *r* = 0.999. The rest of the parameters were estimated from Table 1

In order for the lysogens to displace the nonlysogens, the SPI reaction must occur before the time *T* at which the lysogen population drops to 0. To understand how this time *T* depends on *g* − *r* and *L*_0_, we analytically calculated the latest time an SPI reaction must occur by solving the LUV model (equations 15 - 5) with *σ* = 0 for *L*(*t*). The result is given as in equation 36.

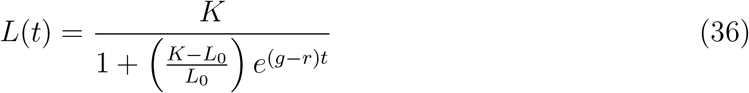

The latest time *T* that an SPI reaction can occur and still allow the lysogens to displace the nonlysogens can be calculated by setting *L*(*T*) = 1 and solving for *T*. We set *L*(*T*) to 1 because by time *T* there is only 1 lysogen left which can undergo SPI. The approximation in 37 is quite accurate since *K* is many orders of magnitude greater than 1, as is *K/L*_0_.

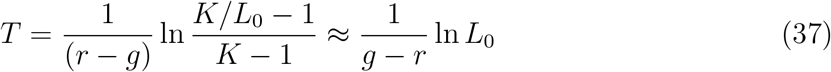

If an SPI reaction occurs by time *T*, then the lysogens will displace the nonlysogens. This establishes some additional constraints on the value of *σ* necessary for the lysogens to displace the nonlysogens, namely that if there is a lysogenic growth cost then the SPI rate cannot be arbitrarily low in a stochastic setting. In Figure 5 we show the *T* value at different values of *r* − *g* and *L*_0_. We see that *T* gradually increases with *L*_0_, implying that higher initial lysogen count gives the lysogens more time to undergo an SPI event. However, we see that as *r* becomes closer to *g* the value of *T* increases dramatically. This means that the lysogens have a much longer time to initiate an SPI event when there is a small lysogenic growth rate cost. If the lysogenic growth-cost is arbitrarily small, then *r* ≈ *g* → *T* → ∞, implying that eventually an SPI reaction will occur and the lysogens will displace the nonlysogens with probability 1.

**Figure 5:**
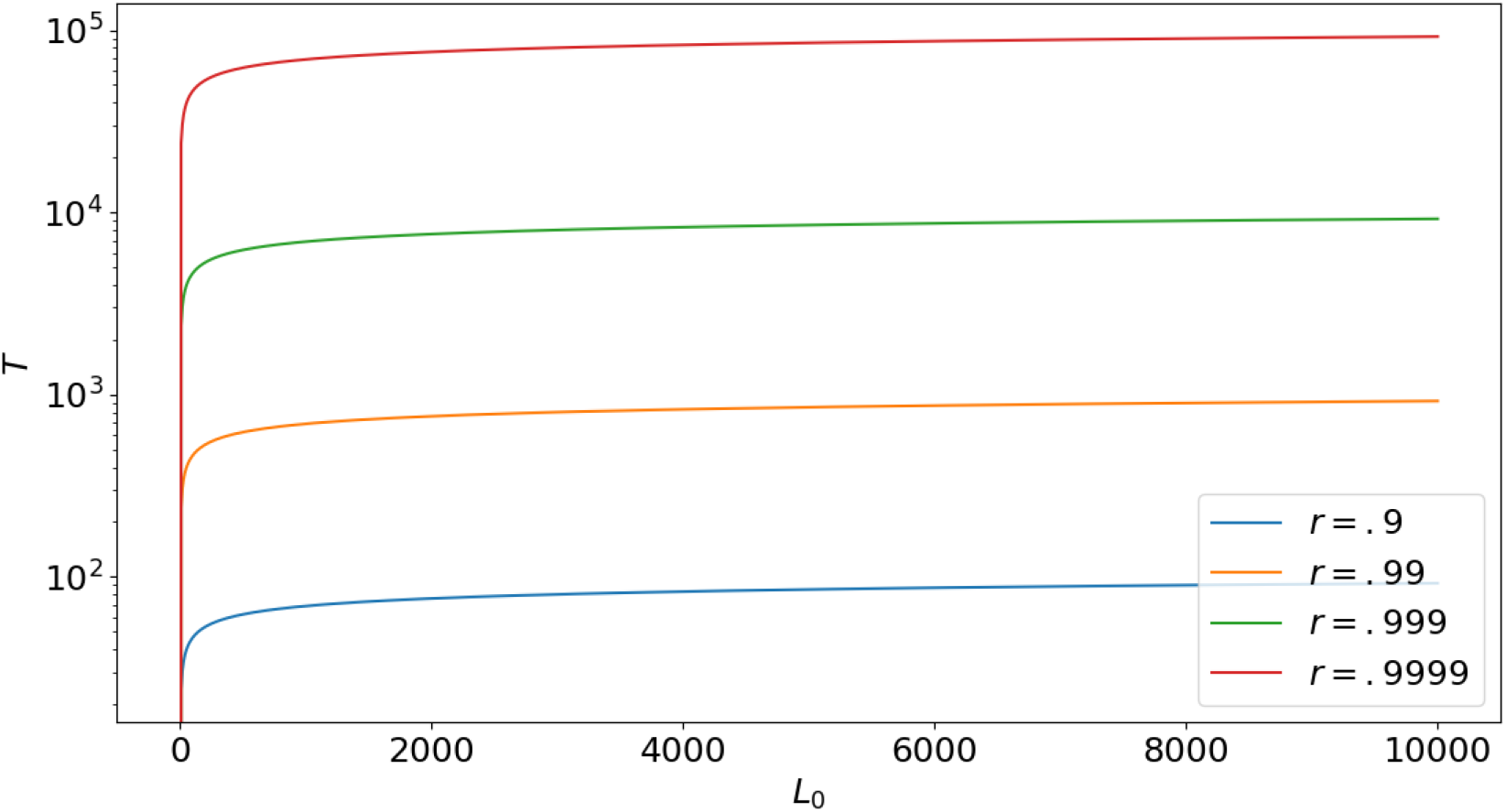
The *T* value calculated at different values of *r* and *L*_0_ using *K* = 10^6^.

To understand how the probability, Ψ, that the lysogens will displace the nonlysogens depends on general parameter values (e.g. not just when *r* ≈ *g*), we calculated the probability that at least 1 SPI reaction occurs within time *T* by modeling our stochastic simulation analytically. First, we split up the time interval 0 ≤ *t* ≤ *T* into small subintervals denoted by Δ*t_k_* of fixed size Δ*t* of which there are *N* = *T*/Δ*t*. Within each of these intervals, the number of lysogens is denoted by *L_k_*. The probability of at least 1 SPI reaction over 0 ≤ *t* ≤ *T* is equal to 1 minus the probability that 0 SPI reactions take place over 0 ≤ *t* ≤ *T*. This is equivalent to 1 minus the probability that 0 SPI reactions occur within all *N* = *T*/Δ*t* subintervals Δ*t_k_*. In each subinterval, the probability of 0 SPI reactions is [1 − *σ*Δ*t*]*^L_k_^*. The total number of subintervals is *N* = *T*/Δ*t*, which implies that the probability of 0 SPI reactions over the entire interval 0 ≤ *t* ≤ *T* is 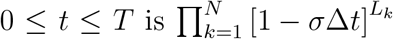. This can be rewritten as [1 − *σ*Δ*t*]^Σ*L_k_*^ where the sum Σ ranges over the *L_k_* from *k* =1 to *k* = *N* = *T*/Δ*t*. Using 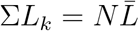 with 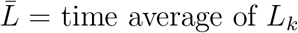, we find that 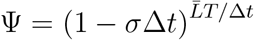. As Δ*t* → 0 one can easily show that Ψ is given by equation 38.

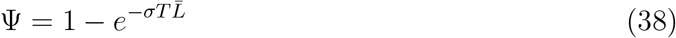

Since we know *L*(*t*) over time *T* from equation 36, we can calculate 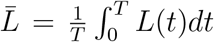. Applying straightforward integration techniques, we find that 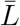 is given by equation 39.

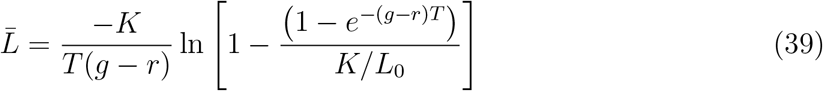

We can then determine a final expression for Ψ by combining equations 38, 37, and 39 to derive equation 40.

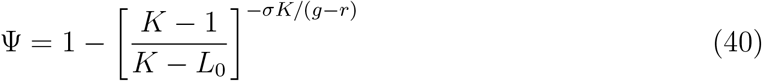

Our expression for Ψ matches the results of our stochastic simulation remarkably well as shown in Figure 6 using the same parameter values from Table 1. From this expression it is clear that the probability that the lysogens displace the nonlysogens increases with *L*_0_, *σ*, and *K*. There is also a strong dependence on *g* − *r*. If the lysogenic-growth cost is very small then *g* − *r* → 0 and, as a result, Ψ → 1 − 0=1. Overall, the lysogens can maximize their probability of displacing nonlysogens by maximizing the quantity *σ*/(*g* − *r*). The value of *σ* can be arbitrarily small so long as the lysogenic growth-cost (e.g. *g* − *r*) is correspondingly small (which it is expected to be). These insights from our analytical models agree with our stochastic simulations. It is worth noting that when *r* ≈ *g* the simulations are extremely time consuming, especially at low *L*_0_, because both the lysogen and nonlysogen population remains approximately constant in time until finally an SPI event inevitably occurs since *σ* > 0. However, from our analytical results we can immediately see the effect of various parameters on *P* without simulation. We take advantage of this by using Ψ to calculate the probability the lysogens displace the nonlysogens over a range of *L*_0_ and *σ* values for *g* − *r* = 10^−5^. Using the stochastic LUV model for the same calculation would have taken 10s of hours or longer.

**Figure 6:**
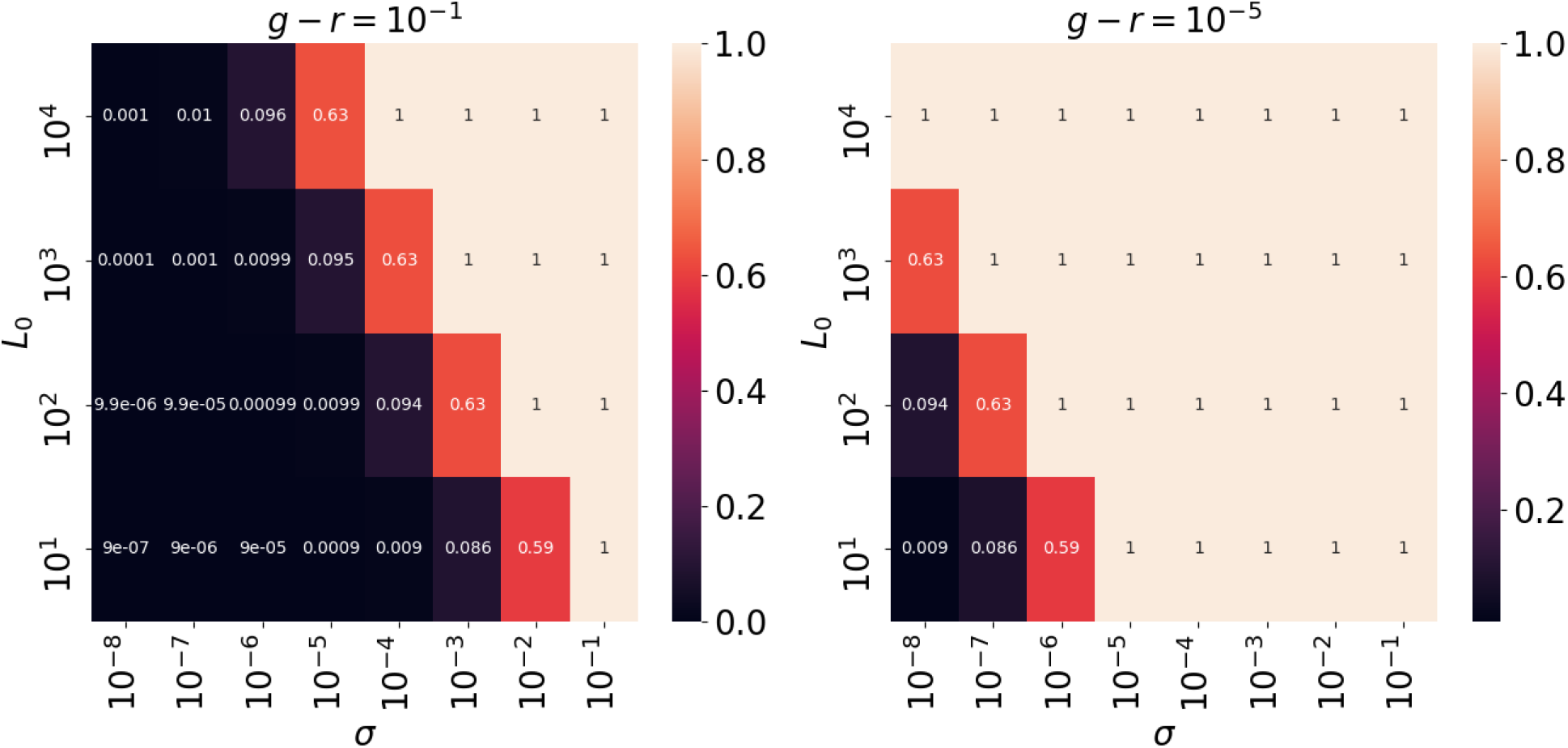
Probability that the lysogens displace the nonlysogens using the Ψ function. Left panel matches with stochastic simulation results in Figure 4, and right panel shows results when *g* − *r* is very small (e.g. 10^−5^) the lysogens are more likely to displace the nonlysogens for any *L*_0_ and *σ*.

To understand how the stochastic evolutionary dynamics occurs in the case that there is a single initial lysogen, we created a separate model because our prior models don’t adequately capture the case when *L*_0_ = 1. In this case, when *t* = 0 we have that *V* = 0, *U* = *K* − 1, and *L* = 1 so the dynamics of *L* can be described by 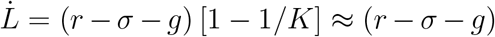 using 1. The approximation holds since 1/K is very small compared to 1. This expression for 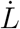 shows that the initial rate of change for L is made of two parts. The *g* − *r* term corresponds to the competitive growth of the lysogens vs the nonlysogens. The second term *σ* corresponds to the SPI reaction. If *g* > *r* then there are only two possibilities for the next reaction, namely i) either the nonlysogens will displace the nonlysogens by outgrowing them or ii) an SPI event will occur and the lysogens will displace the nonlysogens. The time to the next reaction in typical Gillespie fashion is given by *τ* = − ln [1 − *R*] /(*g* − *r* + *σ*) with *r* as a random number from a uniform distribution ranging from 0 to 1. The probability that the next reaction at time *τ* is the SPI reaction is simply given by the proportion *σ*/(*g* − *r* + *σ*) and can be written as in equation 41.

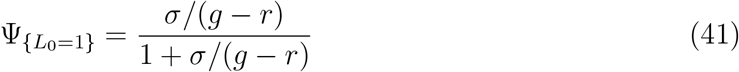

From equation 41 we clearly see that in the case *L*_0_ = 1 the probability the lysogens displace the nonlysogens approaches 1 as *r* → *g*. Thus, at little to no lysogenic growth-cost, the lysogens can displace the nonlysogens with high probability even when there is only 1 initial lysogen. As in equation 40, the higher the ratio *σ*/(*g* − *r*) is, the more likely the lysogens displace the nonlysogens. It is easy to show that Ψ_{_*L*0_=1}_ ≥ 0.5 if *σ* ≥ *g* − *r*. Thus, with *σ* = 10^−5^ the lysogens have a 50% chance of winning if the lysogenic growth-cost is on the order of 10^−5^. If *g* = *r* then in the model both *L* and *U* remain at their initial population sizes until an SPI reaction fires, and so the lysogens win with probability Ψ = 1 in this case. This is easily shown by letting *g* → *r* and taking a limit of equation 41.

To understand how stochastic factors influence the competition between two lysogen strains, we modified the LUV2 model to be stochastic in the same way as in the LUV model. The insights we gained were similar to the case of the LUV model with one extra detail. As shown in Figure 7, we see that so long as either one of the lysogens undergoes SPI then *L*_2_ will always displace *L*_1_ in the long run. This is because when the SPI event initially occurs, the nonlysogens start to be killed off, causing empty space to appear for both *L*_1_ and *L*_2_ to both grow into. Once both *L*_1_ and *L*_2_ fill this empty space and the nonlysogens disappear, it is simply a competition between only *L*_1_ and *L*_2_. From this point onward it is clear that *L*_2_ has the advantage because its effective growth rate *r* − *σ*_2_ is slightly greater than the effective growth rate *r* − *σ*_1_ of *L*_1_ (since *σ*_2_ < *σ*_1_). The *V*_1_ curve jumps up and down a bit towards the end because the stochasticity of SPI reactions becomes more apparent as *L*_1_ drops to lower levels. Notice that *L*_1_ underwent an SPI reaction very early on (marked by the increase in *V*_1_) while *L*_2_’s first SPI event occured much later on at ≈ 10 generations. Even though *L*_2_ was late to SPI, it still wins in the long run since its effective growth rate is higher.

**Figure 7:**
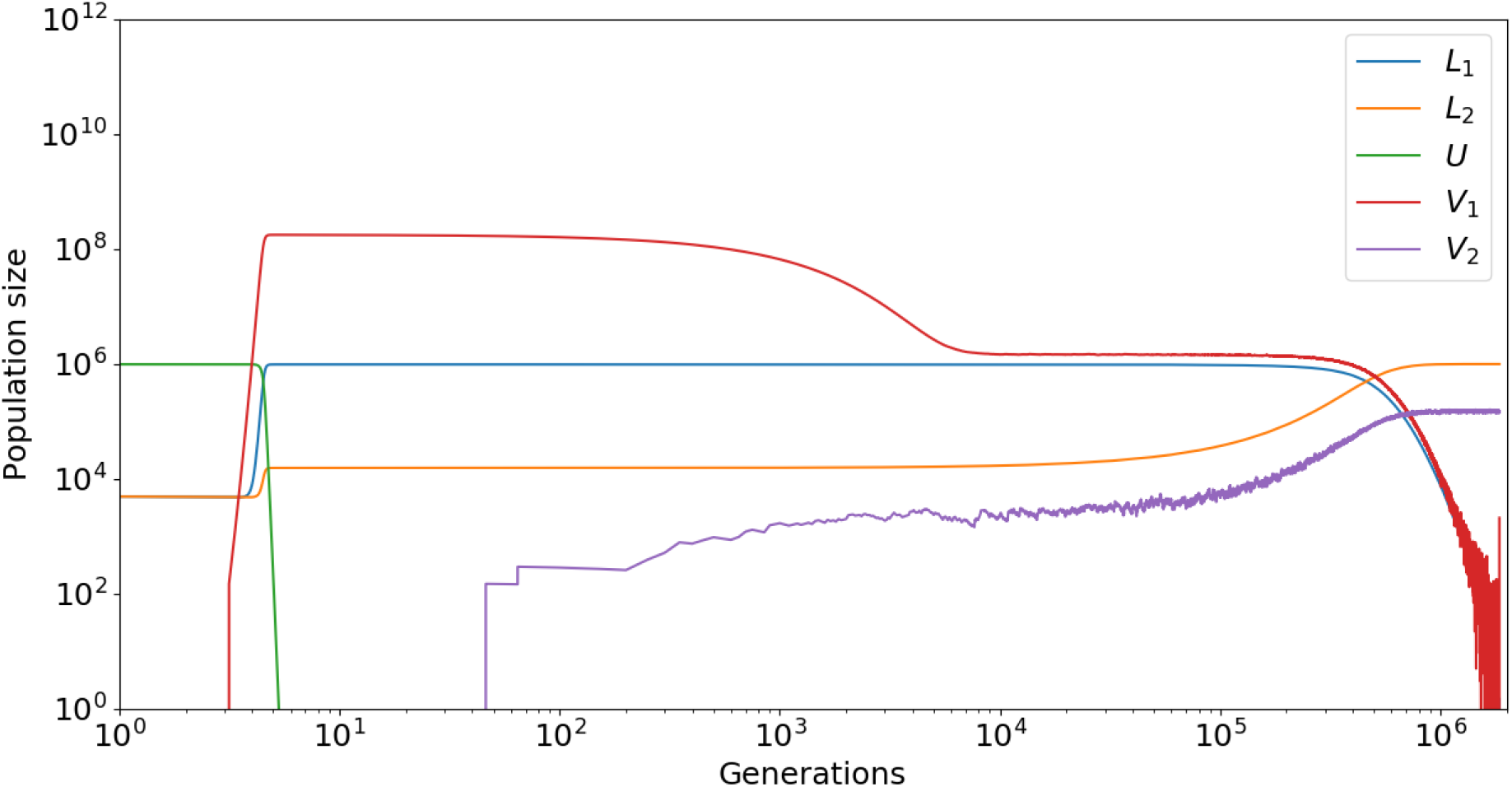
A sample simulation from the stochastic LUV2 model. Early on *L*_1_ manages to have an SPI reaction, causing *V*_1_ to rise and *U* to be killed off. During this, *L*_2_ begins to grow since there is now free space to do so. It does not have its own SPI event until about 10 generations. Eventually, the relatively higher SPI rate of *L*_1_ causes it to slowly but surely drop to 0 as *L*_2_ rises towards the carrying capacity *K* = 10^6^. This simulation was run with *r* = 0.99, *σ*_1_ = 10^−5^, and *σ*_2_ = 10^−6^. We observed the same qualitative behavior for other parameter values.

## 7 Discussion

In this work we explore how SPI influences the natural selection among lysogens and nonlysogens, and also between lysogen variants with different SPI rates. We find that there exists a lower bound *σ_LB_* ≈ 10^−6^ (or smaller) above which lysogens are expected to displace the nonlysogens for general parameter values. Furthermore, we find that lower SPI rates are naturally selected, because although higher SPI rates confer an early advantage they are costly to growth in the long run. These results collectively show that natural selection should push the SPI rate down towards *σ_LB_*, and that this can be viewed as an optimal SPI rate in terms of natural selection. It is plausible that there does exist a lysogenic growth-cost (i.e. *g* − *r* > 0) but it is very likely that this growth-cost is very small such that *g* − *r* ≈ 10^−*n*^ for some large integer *n*. Thus, there likely is a lower bound given by *σ_LB_* that is strictly greater than 0, but it is likely to be very small.

If stochasticity plays a major role in the dynamics, then our same conclusions hold in a probabilistic sense, with lysogenic fixation becoming increasingly probable at higher SPI rates. Thus, stochastic effects on natural selection would tend to favor larger SPI rates. In the stochastic setting, however, the SPI rate is scaled by the lysogenic growth-cost, implying that at very low lysogenic growth-cost the SPI rate can remain arbitrarily low while still maintaining a high probability of lysogenic fixation. In the extreme case that there is only 1 initial lysogens, the probability they will fix in the population is ≈ 1 if the lysogenic growth-cost is on the order of 10^−6^ at the naturally occuring SPI rate *σ*\* ≈ 10^−6^.

Our estimate on the lower bound of the SPI rate is on the order of 10^−6^ or 10^−7^ (or smaller according to *g* − *r*) lysogens per generation, which matches experimental estimates of the *recA*^+^ SPI rate [11, 12, 29]. We show that deterministic evolutionary forces are pushing the SPI rate down towards *σ_LB_*, but stochastic factors tend to push it higher only if *g* − *r* is not small. Otherwise, if *g* − *r* is very small then *σ* can also be very small and still allow natural selection of lysogens over nonlysogens with high probability. Our theory suggests that phage *λ* may be sitting at or near this optimal SPI rate.

Our results demonstrate how mathematical modeling can be effectively used to analyze evolutionary scenarios and explore the consequences of ecological constraints and interactions among competing species. Our theory shows how by sacrificing a little bit of growth rate, a species can gain a competitive advantage over another. This has connections to bacterial persistence, sporulation, and other cell growth strategies in which a small subpopulation switches to a growth-reduced but phenotypically different state. It will be interesting to validate our model further by applying it to other phages and their hosts such as P1, T4, and Mu once all the model’s parameters are measured for these phages.

## 8 Acknowledgements

We acknowledge discussions with the members of the Gabor Balazsi laboratory, as well as helpful comments from Lanying Zeng. This research was supported by National Institutes of Health National Institute of General Medical Sciences (NIH-NIGMS) grant R01GM107597 and administrative supplement 3R01GM107597-02S1 to Lanying Zeng and Gabor Balazsi, as well as by NIH-NIGMS grant R35GM122561 and by a Laufer Center for Physical and Quantitative Biology endowment to Gabor Balazsi. Michael G Cortes received additional support from a W. Burghardt Turner Fellowship, and two Turner Summer Research grants at Stony Brook University.

